# Advancing Remote and Low-Resource Healthcare Delivery with Microbiopsy Skin Sample mRNA Housekeeping Gene Stability Analysis Across Various Temperatures

**DOI:** 10.1101/2024.11.17.624028

**Authors:** Nathalie Nataren, Lucy Gunnell, Tarl W Prow, Miko Yamada

**Author notes:** **Correspondence** Miko Yamada.

## Abstract

In the clinical dermatology setting, conventional skin sampling methods such as punch, shave, or excision biopsy are invasive, requiring local anesthesia and after-procedure care. These methods also necessitate immediate processing and cold storage to preserve high-quality nucleic acid, limiting their use in remote and low-resource regions. The microbiopsy, a minimally invasive device designed for sub-millimetre skin sampling, offers an alternative that may overcome these limitations by providing genetic profiles from the viable epidermis.

We examined skin patient samples (5 microbiopsies, 3 individuals) stored across five conditions temperature conditions expected to be encountered during commercial shipping. RNA was extracted for TaqMan assay Real-Time PCR on the Fluidgim Dynamic Array targeting three housekeeping genes (ACTB, GAPDH, and RPLP0). Housekeeping genes were successfully detected at comparable levels in samples across non-control conditions, contrasting with positive control samples which showed a trend toward lower gene expression detection.

In conclusion, we have established that microbiopsy samples stored in the RNA preservation agent can withstand uncontrolled temperatures likely to be encountered during commercial shipping. This could have a significant impact on enabling access to molecular diagnostic testing in an era of increasing personalized medicine and to address the long-standing need to improve remote healthcare delivery.

## Introduction

In the clinical dermatology setting, skin sampling is a fundamental technique for diagnosing and monitoring a range of conditions, from benign lesions to malignancies. Conventional skin sampling consists of shave, punch, or excision biopsy. These well-established techniques, while effective, come with limitations which may impact patient outcomes or practicality in clinical settings. The shave biopsy, which is typically performed using a flexible blade, is the most common due to minimal wound care requirements. Shave biopsy typically yields a flat, thin (<1mm) skin specimen consisting of the epidermis and upper dermis. A punch biopsy captures a cylindrical skin specimen with a diameter of 2-8 mm, to a depth involving the dermis and underlying subcutaneous layer^1^ which can be stored in an RNA preservation agent and frozen at -80°C prior to molecular testing.^2^ Conventional biopsies require local anaesthesia and suturing, which can lead to post-procedure complications such as pain, scarring of cosmetically sensitive areas, and infection.^3^ Tape stripping is an alternative sampling technique which consists of collecting layers of the stratum corneum with an adhesive tape for analysis of mainly DNA and proteins.^4^ Stratum corneum has residual RNA at extremely low concentration (picogram). Compared to the conventional biopsy, tape stripping is a simpler procedure which can be conducted with minimal resources. An example of this technology is the pigmented lesion assay (PLA) which was developed by DermTech Inc (La Jolla, California) to detect genes *PRAME* and *LINC00518* to aid in the identification of melanoma.^5^ The PLA consists of a small adhesive tape that is applied over the suspicious lesion on the patient’s skin, and is stored and sent to DermTech for RNA extraction and subsequent gene quantification through Real Time-PCR. Limitations to tape stripping sampling such as the PLA are that they cannot collect viable epidermis and recent studies report a high proportion of non-actionable results and discrepancies between PLA and histopathology in cases that were biopsied.^6, 7^ With the limitations of current skin sampling techniques, there is a need for a minimally invasive technology capable of sampling all skin regions, can be used with minimal training and which preserves the sample for transit to enable molecular testing for patients in remote and resource-limited settings.

The skin microbiopsy device represents a significant advancement in dermatological practice, addressing the limitations of traditional methods, whilst providing a minimally invasive, yet effective method for obtaining skin samples. The device uses microneedle technology, capable of sub-millimetre skin sampling. The microbiopsy chamber has dimensions of 400 × 50 × 150 μm and captures approximately 400 cells (theoretical sample size of 0.003 mm^3^). The advent of microsampling is driven by improvements in molecular detection techniques, which increasingly require smaller sample sizes for accurate analysis.^8^ The miniaturisation of sampling devices is anticipated to broaden sampling opportunities by moving the sampling closer to point-of-care. Another key advantage of the microbiopsy is the ability to capture skin samples sans the requirement of local anaesthetics or suture and has been used successfully in patients with fragile skin.^9^ The innovative design of the microbiopsy offers a minimally invasive method of sampling viable skin with nominal post-procedure care requirements, positioning the device as a solution for the delivery of molecular testing in resource-limited settings.

The device’s utility has been demonstrated in several clinical settings, including the Munich Atopy Prediction Study (MAPS), which investigates clinical and molecular risk factors for atopic dermatitis in early childhood.^10^ In MAPS, microbiopsy has enabled the longitudinal sampling of skin in a paediatric population, where skin biopsies are traditionally performed less commonly than in adults.^11^ Conventional techniques provoke apprehension in parents or caregivers^12^ and cause patient discomfort, including pain, anxiety and fear, as well as the potential for visible scarring.^3^ These factors may lead to reluctance particularly in undergoing repeat biopsies and may explain why few longitudinal studies of skin disorders using punch biopsy methods are available. The ability to perform repeated sampling with minimal invasiveness or discomfort is a key advantage of the microbiopsy, as it facilitates the monitoring of disease progression or treatment efficacy over time. This case underscores the devices utility in both longitudinal and paediatric settings where patient comfort and adherence are critical. Microbiopsy, thereby has the potential to provide valuable insights into chronic conditions, even in vulnerable populations.

Moreover, the microbiopsy’s design and functionality show significant promise for enhancing dermatological care in the context of global health. The device’s portability, ease of use, and low production cost ensure it is well-suited for deployment in resource-limited environments. In such settings, the ability to perform safe and effective skin sampling may be constrained by infrastructure.^13^

For example, the requirements for conventional biopsy such as sterile environments, anaesthesia, and suturing, can be of low availability.^14^ Meanwhile, the microbiopsy has minimal resource requirements and straightforward application, therefore can be employed by healthcare providers with limited training. This expands its scope to remote and underserved regions, and consequently may contribute to improving outcomes in regions where traditional biopsy methods are not supported.^15^

Furthermore, it is the microbiopsy’s ability to capture viable epidermis which positions it as a critical tool in the domain of personalised medicine. Current methodologies lack the precision necessary for obtaining high-quality viable epidermis, a gap the microbiopsy effectively fills. Conditions localised to the epidermis, such as psoriasis, actinic keratosis, and atopic dermatitis, could benefit greatly from this device. The microbiopsy enables the collection of samples that accurately reflect localized gene expression changes, by selectively selecting the epidermis, which is crucial for developing targeted and effective treatments. To be suitable for molecular testing, RNA extracted from microbiopsy obtained samples must remain stable in varying storage conditions. The extracted RNA also needs to be evaluated for reliable molecular assay performance across conditions during likely transport scenarios likely to be encountered in remote healthcare settings. The aim of the current study was to evaluate whether microbiopsy samples can be effectively preserved under likely commercial shipping conditions and to assess the impact of temperature fluctuations on the quality and stability of extracted RNA for accurate gene expression analysis.

## Methods

In this study, we examined samples stored across five conditions. For the positive control, samples were stored at -20°C for one week in RNA preservation agent, this being common practice for tissue preservation prior to nucleic acid extraction. Samples were then stored at 25°C for one week in RNA preservation agent, the most likely average shipping condition, 37 °C for one day in RNA preservation agent which is the maximum temperature at which RNA preservation agent guidelines state samples will remain intact, and at 47°C (+10°C the maximum RNA preservation agent storage conditions) for one day in RNA preservation agent as a negative control, for which we anticipated a higher level of RNA degradation. Samples were also field tested at an uncontrolled ambient temperature in RNA preservation agent (4,000 km interstate transit by air). The expression levels of housekeeping genes with increasing stability in skin (*RPLP0* > *GAPDH* > *ACTB*) were examined.^16, 17^ Three volunteers participated in the study as per approval from the University of South Australia Human Research Ethics Committee (UniSA HREC # 200607). Five microbiopsy samples were taken from the inner forearm of each participant. The microneedle component of the microbiopsy was immediately removed and stored in RNA preservation agent. Post storage, we extracted total mRNA from individual microbiopsy samples (PicoPure TM,Thermo Fisher Scientific) and performed Real-Time PCR for *ACTB, GAPDH* and *RPLP0* gene expression using 20X TaqMan assays (Thermo Fisher Scientific) on the Fluidigm Dynamic Array Gene Expression system.

## Results

In this study, we assessed the integrity of RNA extracted from microbiopsy obtained skin samples that were stored under various temperature conditions, with the aim to determine the reliability of gene expression analysis for potential use in remote diagnostics. The analysis of gene expression stability in microbiopsy samples revealed no statistically significant differences in the expression levels of *ACTB, GAPDH* and *RPLP0* across the five storage conditions, confirming the robustness of RNA preservation under these conditions. (Figure 1). Positive control samples displayed a trend to lower-than-expected expression across all genes, as indicated by the higher mean Ct values. The positive control samples were not snap frozen prior to storage at -20°C, which may have resulted in RNA degradation by freeze-thawing, which would be more evident in a small sample of approximately 400 cells. The negative control samples displayed moderately decreased gene expression compared to other samples as indicated by the Ct values, but it is important to note that there was an increase in non-detects, particularly for the two less stable genes *ACTB* and *GAPDH* (Table 1).

**Table 1:** House-keeping gene qRT-PCR non-detects amongst biological replicates.

| Gene | -20°C, 1 week (+ ctrl) | 47°C, 1 day (- ctrl) | 25°C, 1 week | 37°C, 1 day | Field test |
| --- | --- | --- | --- | --- | --- |
| *ACTB* | 0 | 2 | 0 | 0 | 0 |
| *GAPDH* | 0 | 2 | 0 | 0 | 0 |
| *RPLP0* | 0 | 1 | 0 | 0 | 0 |
| *Abbreviations: ND = non-detects, ctrl = control* |  |  |  |  |  |

**Figure 1:**
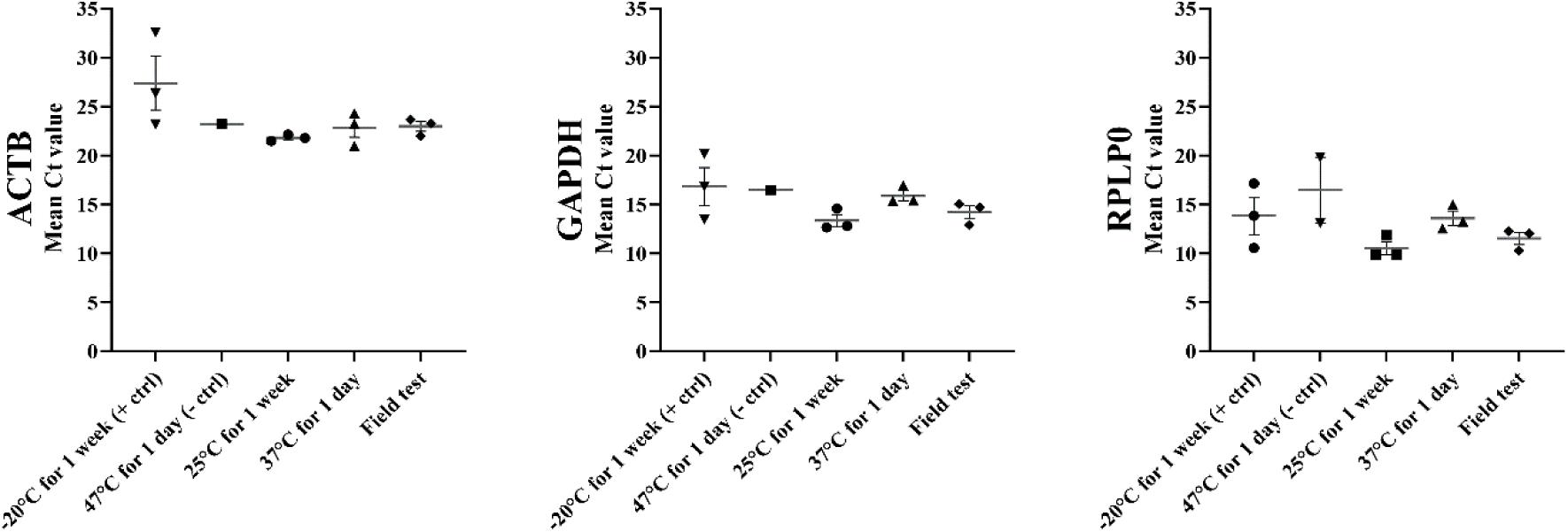
Housekeeping gene expression levels across storage conditions. Gene expression levels at -20°C for 1 week, 25°C for 1 week, 37°C for 1 day, 47°C for 1 day and field test conditions for ACTB, GAPDH and RPLP0. N = 3 individuals per group, each data point is the mean of two technical replicates, expressed as mean ± SEM.

These results support the hypothesis that there is greater RNA degradation likely due to the harsh temperature conditions of the negative control. Microbiopsy samples stored in RNA preservation agent at 25°C degrees displayed the highest expression across all genes. Housekeeping genes in samples for all individuals stored at 37°C degrees were detectable by RT-PCR. The gene expression levels where higher compared to other samples except for the 25°C for 1 week. Notably, housekeeping genes where detectable within the field tested microbiopsy samples, despite the inherent variability of ambient temperatures for this condition, with only the samples stored at 25°C degrees having lower Ct values.

## Discussion

The conventional biopsy methods in dermatology, such as shave and punch biopsies, have been the standard for obtaining skin samples, but generally more invasive and require post-procedure care. Tape stripping is an alternative technique used in dermatology, but its application is limited as it cannot be used on mucosal and acral regions or regions where hair or blood are present due to the loss of traction with the skin The microbiopsy is a less invasive technique, which can be used on all skin regions, and eliminates the need for local anaesthesia or sutures.

Previous studies assessed the optimal conditions for extraction of high-quality RNA from conventional biopsies, finding that skin samples require some form of immediate cold storage or processing prior to RNA extraction, such as tissue sectioning followed by bead homogenisation with cold lysis buffer^18^, immersion in beta-mercaptoethanol supplemented lysis buffer and freezing on dry ice^19^, liquid nitrogen immersion and cryosectioning prior to -20ºC storage in RNALater™,^20^ and Allprotect™ immersion prior to ^18-20^ -20ºC storage and subsequent sectioning.^21^ Whilst these studies provide robust methods for extracting high-quality RNA from conventional biopsy or surgical excision, all of these previous studies immediately placed samples in cold storage. This study assessed whether sufficient RNA can be extracted to perform molecular assays from microbiopsy samples that are stored and transported in uncontrolled without immediate processing or cold storage. The experiment’s design, which included different temperature conditions likely encountered during commercial shipping, addresses a critical aspect of sample transportation and storage of samples, particularly from remote regions. The use of RNA preservation agent as a storage medium is a key focus, as it’s commonly used for preserving samples for molecular testing.

The results are particularly intriguing, as they suggest that microbiopsy samples remain stable in RNA preservation agent for housekeeping gene detection under a range of temperatures, including those likely to be encountered during shipping. The lack of significant difference in gene expression across various storage conditions, including the optimal and extreme temperatures, and the ability to detect more labile RNA transcripts, are promising indications of the method’s robustness. The unexpected findings in the positive control samples, which displayed a trend towards lower gene expression than anticipated, raise questions about the potential effects of storage conditions on small sample sizes. This aspect highlights the need for careful consideration of sample handling, especially in the context of small samples like those obtained from microbiopsy. Furthermore, the study has implications for remote molecular monitoring and potentially broadens the accessibility of molecular testing, especially in settings where immediate sample processing is not feasible. Previous studies that have included non-cold storage conditions for samples stored in an RNA preservation also found that the resulting RNA was of good quality and could be used for molecular testing.^22, 23^ However unlike the current microbiopsy study, the tissues in these previous studies underwent some form of preprocessing such as the cutting uterine tissue into <2 mm sections and homogenisation of skin punch biopsies. The gene expression results from the current study demonstrates that microbiopsy obtained samples do not require any additional processing prior to storage and transportation in RNA preservation agent, eliminating a step in the diagnostic workflow which is important in resource limited settings. By demonstrating that microbiopsy samples can be reliably stored and transported at varying temperatures without significant degradation, the study affords new possibilities for skin disease diagnosis and monitoring, particularly in remote or resource-limited settings.

As with any sampling method there are potential limitations or challenges associated with the practical application of the microbiopsy in remote settings. Whilst sampling with the microbiopsy is minimally invasive and could be potentially self-administered, in the context of a suspected malignant lesion, a physician trained in use of the microbiopsy is best placed to ensure proper triaging and correct sampling of smaller suspected malignant lesion. Given that the entirety of a suspicious lesion is not removed by the microbiopsy in contrast to a punch biopsy, the anatomical position and appearance of the lesion on the patient’s body will need to be tracked for the purposes of future treatment. A foreseeable trade off with taking such a small sample is the potential to ‘miss’ the lesion during sampling and extract non-lesional tissue instead. This could be readily mitigated by taking a few microbiopsy samples and pooling them, as was done in this study. A previous study using the microbiopsy demonstrated that three pooled microbiopsy samples was sufficient to detect changes in known melanoma gene expression biomarkers to distinguish a superficial melanoma from a pigmented Basal Cell Carcinoma.^24^ Future translational research should aim to establish clinically validated guidelines regarding the sampling workflow and minimum number of microbiopsy samples required to guard against false negative diagnoses due to sampling errors.

Given the smaller sample size of this study, future work would look at replicating these findings in a larger patient cohort and extending the study to other conditions, such as long-term storage at -80ºC and storage with other preservation and processing agents such as Qiagen lysis buffers, Allprotect™ and even storage without a preservation agent, to expand the applicability of the microbiopsy across diagnostic settings. The trend towards lower gene expression detection in the microbiopsy samples stored at -20ºC in RNA preservation agent™ for 1 week also bears further investigation to determine whether gene detection is improved with liquid nitrogen freezing of microbiopsy samples prior to RNA preservation agent storage.

In conclusion, this study represents a significant step forward in dermatological sample stability for transportation, offering a more patient-friendly, minimally invasive option while ensuring sample integrity for remote molecular testing. The data from this study supports the hypothesis that microbiopsy skin samples in an RNA preservation solution can remain stable under a wider range of storage temperatures without further tissue pre-processing, which could enable skin cancer diagnostics or research in remote and low-resources regions.

